# Topology recapitulates ontogeny of dendritic arbors

**DOI:** 10.1101/2023.02.27.530331

**Authors:** Maijia Liao, Alex D. Bird, Hermann Cuntz, Jonathon Howard

**Affiliations:** Department of Molecular Biophysics & Biochemistry, Yale University, New Haven, USA; Ernst Strüngmann Institute (ESI) for Neuroscience in Cooperation with Max Planck Society, Frankfurt am Main, Germany

**Keywords:** topology, dendrite, morphology, power law

## Abstract

Branching of dendrites and axons allows neurons to make synaptic contacts with large numbers of other neurons, facilitating the high connectivity of the nervous system. Neurons have geometric properties, such as the lengths and diameters of their branches, that change systematically throughout the arbor in ways that are thought to minimize construction costs and to optimize the transmission of electrical signals and the intracellular transport of materials. In this work, we investigated whether neuronal arbors also have topological properties that reflect the growth and/or functional properties of their dendritic arbors. In our efforts to uncover possible topological rules, we discovered a function that depends only on the topology of bifurcating trees such as dendritic arbors: <u>the tip-support distribution</u>, which is the average number of branches that support *n* dendrite tips. We found that for many, but not all, neurons from a wide range of invertebrate and vertebrate species, <u>the tip-support distribution</u> follows a power law with slopes ranging from -1.4 and -1.8 on a log-log plot. The slope is invariant under iterative trimming of terminal branches and under random ablation of internal branches. We found that power laws with similar slopes emerge from a variety of iterative growth processes including the Galton-Watson (GW) process, where the power-law behavior occurs after the percolation threshold. Through simulation, we show the slope of the power-law increases with the branching probability of a GW process, which corresponds to a more regular tree. Furthermore, the inclusion of postsynaptic spines and other terminal processes on branches causes a characteristic deviation of the <u>tip-support distribution</u> from a power law. Therefore, the tip-support function is a topological property that reflects the underlying branching morphogenesis of dendritic trees.

## Introduction

Branching morphogenesis has fascinated biologists and physicists for decades, because of both its complexity and ubiquity. In the case of neurons, Santiago Ramón y Cajal proposed that morphology is crucial for function (Cajal, 1995). The morphology of a neuron’s dendritic tree defines how the cell receives synaptic inputs from other neurons and how these inputs are integrated to allow signal transmission and computation. For example, observed neurite lengths and branching geometries optimize electrical signaling by minimizing propagation times (Rall, 1964; Jaffe and Carnevale, 1999; Cuntz *et al.*, 2007, 2010, 2012, 2021; Wen and Chklovskii, 2008; Bird and Cuntz, 2016). Neuronal processes also deliver nutrients and energy to support the growth and activity of the cell (Sterling and Laughlin, 2015). The observed distributions of dendrite diameters, another important geometric property of neurons, may optimize the intracellular transport of materials for growth and homeostasis (Williams *et al.*, 2016; Sartori *et al.*, 2020; Liao *et al.*, 2021; Donovan *et al.*, 2022). Thus, geometry constrains the transport of signals and materials through the cells, and leads to the dendrite properties such as length and diameter that scale with the size of the cell (Marshall, 2020). In this paper we asked whether dendrite topology also constrains or reflects neuronal function or growth.

The extreme diversity of the branching patterns of neurons (Markram *et al.*, 2004) has also attracted effort to categorize neuronal morphologies into smaller numbers of distinct classes using topological concepts (Markram *et al.*, 2004; Ascoli *et al.*, 2008; DeFelipe *et al.*, 2013; Gouwens *et al.*, 2020) in addition to geometry. One avenue is topological data analysis (TDA), which is an approach to the analysis of data using techniques from topology to obtain information that is independent of the particular metrics (Carlsson, 2009); this approach can reveal a system’s intrinsic structure and distinguish that structure from other structures and noise (Horak *et al.*, 2009; Giusti *et al.*, 2015; Sizemore *et al.*, 2019). The topological morphology descriptor (TMD), based on TDA, encodes the spatial structure of branched trees as a ‘barcode’ and has been found to be useful for categorizing neurons (Kanari *et al.*, 2018). In addition to TMD, which includes geometric information, there are several other approaches that consider purely topological properties, namely those that are invariant under homeomorphisms of the 2D or 3D space in which the neurons are embedded. These approaches include tree asymmetry (van Pelt *et al.*, 1992; van Pelt *et al.*, 2001), Strahler ordering (Strahler, 1952; Berry and Bradley, 1976; Vormberg *et al.*, 2017), and subtree persistence (Scheele *et al.*, 2017). Because the genetic networks responsible for pattern formation are frequently conserved through evolution, neuronal structures with different sizes are often described by self-similar functions, such as the fractal dimension (Caserta *et al.*, 1990; Marks and Burke, 2007), and one might anticipate that a single set of pattern-formation rules could be used to generate many arbors (Lindenmayer, 1968; Ascoli and Jeffrey, 2000; Wen *et al.*, 2009; Snider *et al.*, 2010). Thus, it is possible that neurons possess scale-free topological properties.

In this study, we imaged and analyzed invertebrate (Jan and Jan, 2010; Scheffer *et al.*, 2020) and vertebrate neurons (NeuroMorpho.Org, Ascoli *et al.*, 2007) and discovered a topological property of dendrites, <u>the tip-support distribution</u>, which is the distribution of the average number of branches that support a given number, *n*, of dendrite tips. For many neurons, <u>the tip-support distribution</u> satisfies a power law, which can be explained quantitatively through a remarkably simple and invariant design principle based on percolation theory. The slope of the power law reflects the bifurcation probability of neuronal dendrites: the higher the bifurcation probability during growth, the higher the log-log slope. The presence of postsynaptic spines on mammalian neurons and branchlets on *Drosophila* class III dendritic arborization (da) sensory neurons, however, leads to a characteristic deviation from the power law suggesting that they rise through a different growth mechanism compared to the primary branches. Thus, the tip-support distribution is a topological property that reflects the underlying branching morphogenesis of dendritic trees.

## Results

### Tip-support distribution follows a power-law for class IV neurons

In our quest to uncover underlying growth mechanisms of neurons, we investigated the topological properties of class IV da neurons in *Drosophila melanogaster* larvae, which are a model system for studying dendrite morphogenesis (Jan and Jan, 2010). We found that <u>the tip-support distribution</u>, a topological property of binary trees, of which dendritic arbors are an example, has a particularly simple form in these cells. The tip-support distribution is the average number (or density), *M*, of branches that support *n* dendrite tips (or leaves). We discovered that the tip-support distribution, *M*(*n*), of class IV cells follows a power law, *M*(*n*)~*n*^-*α*^, with an average exponent *α* of 1.4 (Fig. 1C) with tip numbers range from 1 to 1000, the maximum number of tips at the end of larval development.

**Fig. 1.**
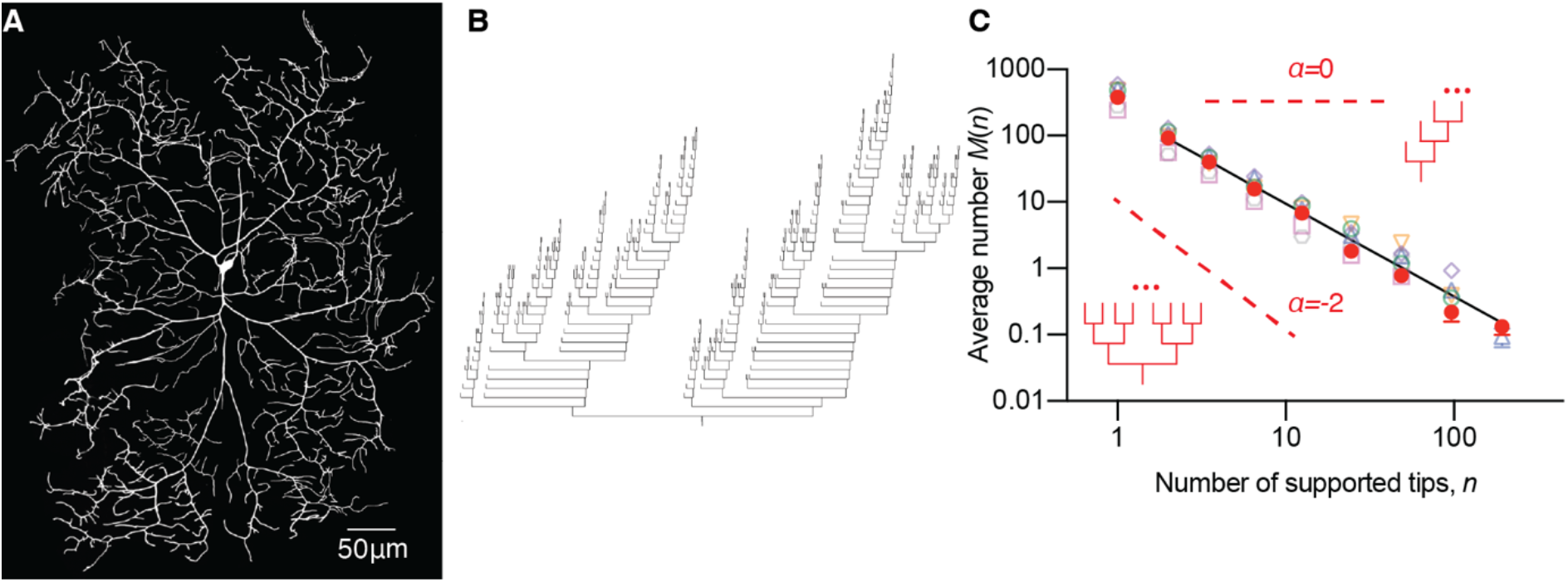
Tip-support distributions for class IV sensory neurons follow a power-law distribution. **A**. Dendritic arbor of a 96hr class IV neuron visualized with a GFP-tagged membrane marker by spinning disk confocal microscopy (see Materials and Methods). **B** Dendrogram of the upper half of the arbor in A. **C**. Tip-support distribution for seven different dorsal class IV neurons from segment A3 to A5 in larvae. The solid symbols correspond to the cell in A and B. The solid black line is the RMA fit to the log-log data with a slope of −1.40 ± 0.04 (mean ± SD). The inserts show a perfect tree (lower left) and an imperfect tree (upper right) with slopes −2 and 0 respectively.

### The perfection index for binary trees

The tip-support function is related to the probability of branching at each node. In a perfect binary tree, namely one in which every branch node has exactly two child nodes, and all leaves are at the same depth (Fig. S1A), the tip-support distribution is a power law with slope −2 (see Methods). This is illustrated with the following example for a tip number *n* = 521 = 2^9^. Then, the number of branches with 1, 2, 3, … tips is *N*(1) = 2^9^, *N*(2) = 2^8^, *N*(3) = 0, *N*(4) = 2^7^, *N*(5) = *N*(6) = *N*(7) = 0, *N*(8) = 2^6^, etc. If we plot the <u>average number</u> of branches *M* against *n* on a log-log axis with logarithmic binning then the power-law slope is −2.0 (Figure 1C inset lower left). The other extreme case is a maximally imperfect binary tree, in which each leaf has a different depth, like a half fishbone (Fig. S1B). In this case, *N*(1) is the total number of leaves and *N*(*n* > 1) = 1. The power-law slope in the log-log plot of *Mvs n* is 0 (*n* > 1) (Figure 1C inset upper right). Based on these empirical properties, we propose that if *M*(*n*) scales linearly with leaf number *n* (on a log-log plot), with power-law slope *-α*, we define the <u>perfection index</u> *β* = *α*/2. Therefore, *β* is 1 for a perfect binary tree (oblique red dashed line in Fig. 1C) and 0 for a maximally imperfect tree (horizontal red dashed line in Fig. 1C). Class IV dendrites are imperfect. The average perfection index of class IV neurons is 0.7, which lies between the two extreme cases.

### The tip-support distribution has a power law over a wide variety of neurons with perfection indices ranging from 0.7-0.9

To test whether the perfection-index concept can be generalized to other neurons, we analyzed Purkinje cells (Fig. 2A) and retinal ganglion cells. To obtain enough statistics for *M*(*n*), we choose neuronal dendrites with more than 60 terminal branches. Fig. 2B shows that the power law holds for Purkinje cells (Rapp *et al.*, 1994; Vetter *et al.*, 2001; Anwar *et al.*, 2014) and retinal ganglion cells (Werginz *et al.*, 2020). The exponent *α* of the power-law ranges from 1.5 to 1.8. The plot of residuals for the example cell types confirmed the goodness of fit (Fig. S2).

**Fig. 2.**
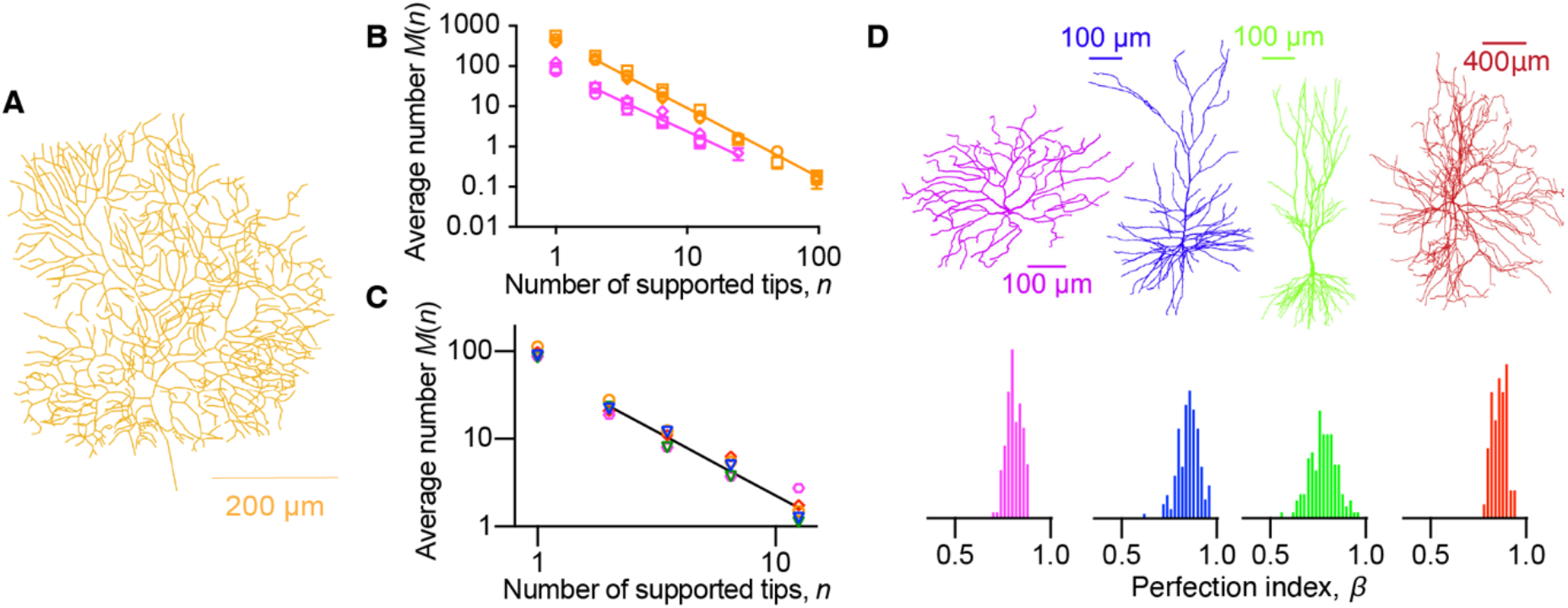
Tip-support distributions and perfection indices for dendritic arbors of vertebrate and invertebrate central neurons. **A.** Skeletonized guinea-pig cerebellar Purkinje cell (Rapp *et al.*, 1994) in which branches have been replaced by their center lines. **B.** Tip-support distributions for Purkinje cells (orange) and retinal ganglion cells (magenta). The RMA slopes are −1.72 (Purkinje cells) and −1.52 (retinal ganglion cells). **C**. Five *Drosophila* T5 cells are shown in Fig. S3. The power-law exponent is 1.46. **D.** A collection of tree perfection indices measured from reconstructed neurons from the morphological database www.neuromorpho.org. The color corresponds to neuronal type: retinal ganglion: 0.80 ± 0.04 (magenta, *n*=130 cells), neocortical pyramidal: 0.85 ± 0.06 (blue, *n*=165 cells), hippocampal pyramidal cells: 0.78 ± 0.07 (green, *n*=131 cells), and motoneurons: 0.86 ± 0.04 (red, *n*=56 cells). All errors are standard deviations.

Next, we tested whether the narrow range of power-law exponents holds across diverse dendrite branching morphologies in a large number of different cell types. In addition to *Drosophila* peripheral, sensory class IV neurons, we also studied T5 cells (Fig. S3) of the adult *Drosophila* central nervous system (Scheffer *et al.*, 2020). The size of these cells remains relatively stable in adulthood, in contrast to class IV cells, which grow throughout larval development. As shown in Fig. 2C, the power law holds with an average exponent of 1.5. We further analyzed the dendritic trees of different cell types from the NeuroMorpho database (Ascoli *et al.*, 2007) including Purkinje cells from the cerebellums of guinea pig, rat, and mouse; spinal motoneurons from rat and cat; retina ganglion cells from mouse, pouched lamprey, and salamander; pyramidal cells in the hippocampus of rat, mouse and guinea pig; pyramidal neurons in the neocortical layers of rat, mouse, cat, monkey, and human. The power law holds for these neurons. The perfection index of individual cells from flies to mammals ranged from 0.7 *(Drosophila* class IV neurons) to 0.9 (cerebellar Purkinje cells), with a mean and SD across all measured neurons of 0.82 ± 0.07 (*N* = 509 cells). Thus, the tip-support distributions follow power laws with similar exponents for a wide variety of vertebrate and invertebrate neurons.

### Perfection Index is invariant under trimming of terminal branches and random ablation of internal branches

When we iteratively trimmed the terminal branches of class IV neurons, the tip-support distribution still followed a power law, with similar exponents to the original neuronal tree, with *α* in the range of 1.4 to 1.5 (Figs. 3A-D). Purkinje cells showed similar behavior (Fig. 3E). When we randomly ablated internal branches of class IV neurons (Figs. 3F and 3G) and Purkinje cells, tip-support distributions also followed power laws (Fig. 3H) with perfection indices remaining invariant (Fig. 3I). Thus, the perfection index of neurons is invariant under such perturbations.

**Fig. 3.**
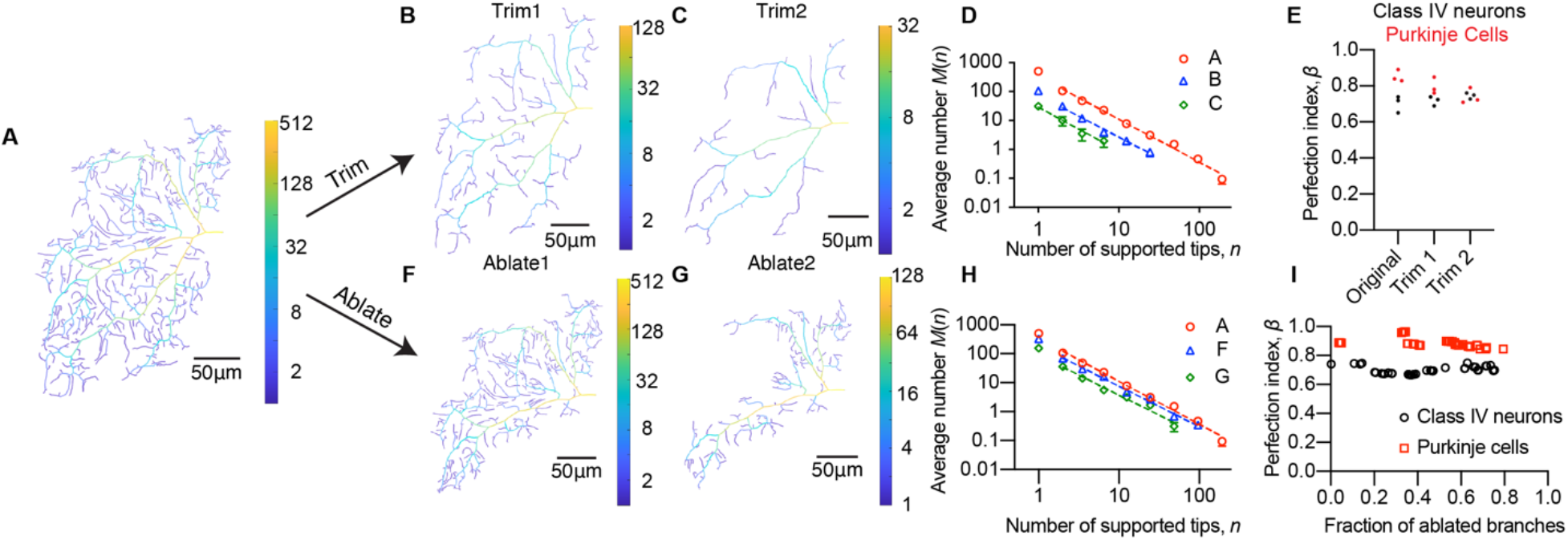
The perfection index is invariant under trimming and ablation. **A.** Part of a class IV arbor with supported tips colored according to the scale at right; **B.** Arbor A after trimming (i.e., removing terminal branches); **C.** Arbor A after trimming a second time; **D.** The power-law slopes are −1.48 (A), −1.45 (B), and −1.52 (C) respectively. **E**. Perfection indices were measured from the original tree and from trimmed trees of class IV neurons and Purkinje cells. **F.** Arbor A after randomly ablating 35% of the whole branches; **G.** Arbor A after randomly trimming 69% of the whole branches. **H.** The power-law slopes are *α* = −1.48 (A), *α* = −1.39 (F) and *α* = −1.38 (G). **I.** Perfection indices from class IV neurons and Purkinje cells following branch ablation.

### Percolation transition associated with the Galton-Watson branching process

To gain insight into why neurons have tip-support distributions that can be described by a power law with a fairly narrow range of exponents, we studied the Galton-Watson (GW) model, a simple stochastic process that generates random bifurcating trees (Peter, 1975). In this model, each branch point (denoted by B) bifurcates into two new branch points with probability *p*, or stops bifurcating to form a tip (denoted by T) with probability 1 – *p*:

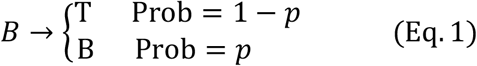

When *p* = 1, the GW model produces a perfect tree, which is deterministic, and when *p* < 1, the GW produces imperfect trees, which are stochastic in the sense that arbors may vary from cell to cell.

The GW process (illustrated in Fig. 4A) typically produces random binary trees (red dashed lines in Fig 4A) that lie in the space of all possible binary trees (bold black lines in Fig. 4A). For a given binary tree, if the bifurcation probability *p* > 0.5 then on average at least one of the terminal branches will continue to bifurcate. The tree size will then be expected to increase with branch order. Otherwise, when *p* < 0.5, the average tree size will decrease with the branch order and growth will almost surely terminate. The critical bifurcation probability *p* = 0.5 therefore marks a qualitative change in the behavior of a branching process, with supercritical (*p* > 0.5) trees able to reach arbitrarily high branch orders. This behaviour is analogous to a percolation transition, which describes the emergence of long-range connectivity in random systems after a certain value known as the percolation threshold is exceeded.

**Fig. 4.**
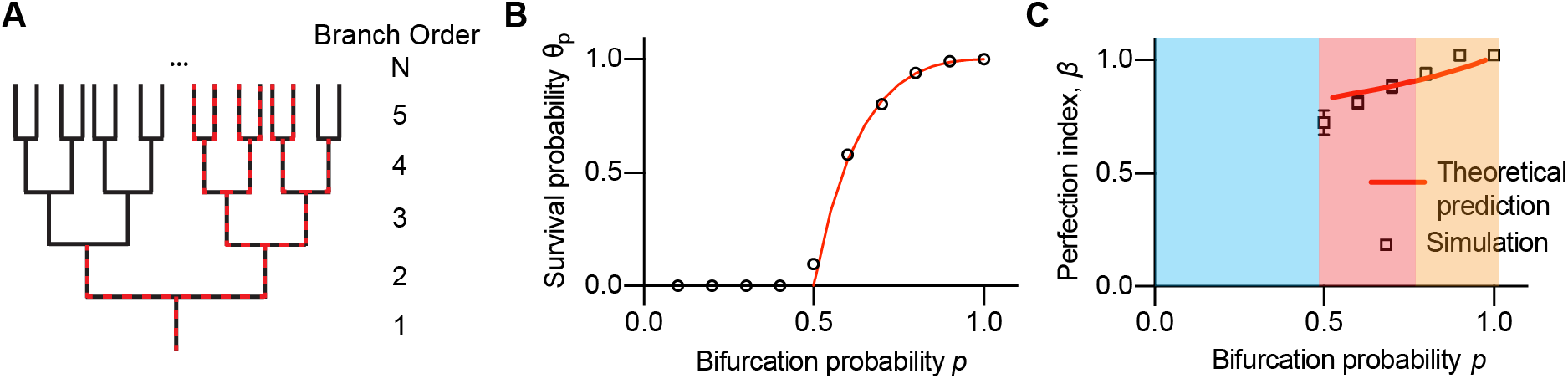
Percolation transition associated with the Galton-Watson branching process. **A.** A sample binary tree (red dashed lines) superimposed on a perfect binary tree (black solid lines). **B.** The survival probability of a Galton-Watson process measured from 1000 simulations (open circles) as a function of the bifurcation probability. The red curve is the analytical formula (Equation (2)). The percolation transition occurs at *p* = 0.5. **C.** Perfection index *β* as a function of branching probability *p*. The theoretical prediction from Eq. (A2) is shown by the red curve. For each branching probability, trees with total leaf number of around 400 were analyzed with 100 simulations for each bifurcation probability. The blue-shaded region corresponds to the region before the percolation transition. The red-shaded region falls within the tree perfection index range observed from a variety of neurons determined (mean - 2SD, mean + 2SD). The orange shaded region is outside the observed range.

To explore the properties of trees around the percolation threshold *p_c_* = 0.5, we defined the survival probability (denoted by *θ*_p_) as the probability of a binary tree reaching arbitrarily high branch orders after branching once. When the bifurcation probability is smaller than the percolation threshold, *θ*_p_ remains zero and trees will almost surely terminate after a finite number of iterations. Above this threshold, *θ*_p_ monotonically increases and reaches 1 when *p* = 1 (Fig. 4B). Simulation results (open black circles) are well fit by the theoretical prediction of *θ*_p_ (red curve) derived in the Appendix:

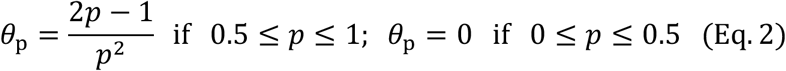

Here *θ*_p_ plays the role of an order parameter, and we can expand it around the percolation threshold *p_c_* = 0.5:

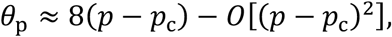

which indicates the critical exponent is 1. Consequently, the stochastic bifurcation process can be described by a continuous phase transition.

We further examined whether the tip-support distribution follows a power law by analytically formulating the stochastic branching process as shown in the Appendix. By solving the equation Eq. (A4), we find that the power law emerges after the percolation threshold with an exponent *α* gradually increasing from 1.6 to 2.0 and *β* = *α*/2 increases from 0.8 to 1 (*p* > 0.5, the red curve in Fig. 4C). Power laws with similar exponents are also observed for Galton-Watson simulations (Fig. S4 and black squares in Fig. 4C). Interestingly, bifurcation probabilities larger than the percolation threshold *p_c_* = 0.5 and smaller than 0.8 (Figure 4C red region) correspond to the perfection indices observed over a wide variety of reconstructed neurons.

### Variation in bifurcation probability does not change the power-law behavior of the tip-support distribution

Neurons come in a variety of shapes (Markram *et al.*, 2004, 2015). Neuronal arbors are shaped by many processes and constraints including stochastic branching and growth (Shree et al. 2022), morphogen gradients (Lin *et al.*, 2021), self-avoidance (Grueber and Sagasti, 2010) and other environmental effects (Pannese, 2015), and available space (Niven and Farris, 2012). To generate a finite tree, the bifurcation probability *p* must go below 0.5 at higher branch order. This could happen in two ways: (a) there is an abrupt decrease in *p* (to zero) at some time or branch order and (b) *p* might decrease to a value below the critical value of *p_c_* = 0.5, so that the tree slowly stops growing as found in class IV neurons (Fig. 5C). In this section, we will investigate how changing the bifurcation probability changes the power-law behavior of the tip-support distribution.

**Fig. 5.**
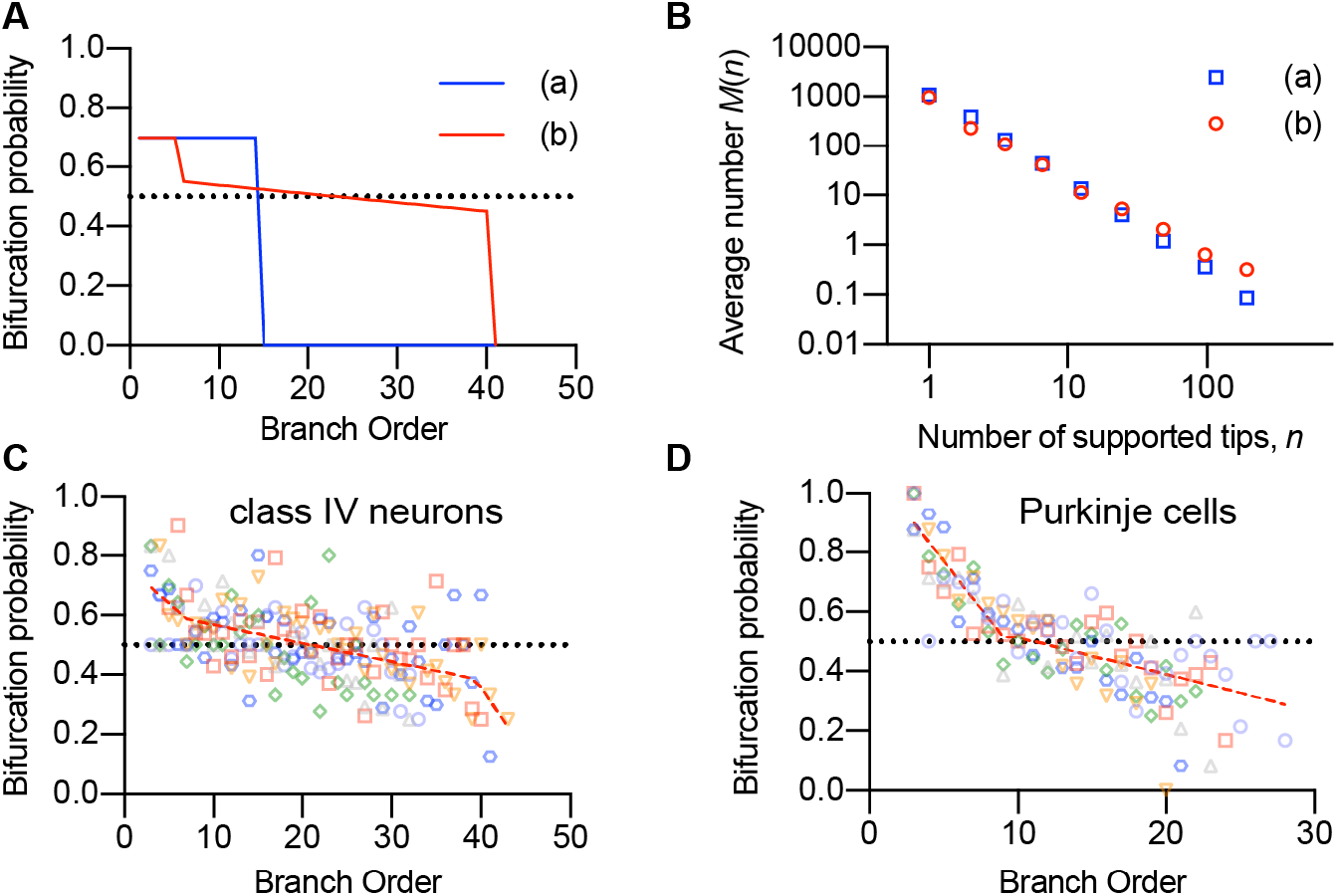
Perturbation of the bifurcation probability changes the log-log slope of the tip-support distribution. **A.** Two bifurcation probabilities functions. The blue curve shows an abrupt decrease of bifurcation probability from 0.7 to 0 at branch order 15. The red dashed curve shows a decrease from 0.7 to 0.55 at branch order 6 and a gradual change from 0.55 to 0.45 from branch order 7 to 40, followed by an abrupt change to 0. **B.** *M(n)* vs. *n.* for the two cases in B show power-law distributions, though the slopes differ. The exponent in (a) is *α* = 1.79 and in (b) is *α* = 1.52. **C** and **D.** Bifurcation probabilities as a function of branch order for class IV neurons and Purkinje cells respectively. In **C** and **D**, six neurons were analyzed with colors representing different animals. Branch orders with more than six branches were chosen and analyzed for each neuron. The black dotted line shows a bifurcation probability of 0.5. A linear piecewise fit (dashed red lines) is overlaid.

We performed simulations using the branch-order dependent bifurcation probabilities shown in Fig. 5A. In both cases, the tip-support distribution *M*(*n*) vs. *n* follows a power law as shown in Fig. 5B. Binary trees grown with higher bifurcation probabilities at lower branch order have higher perfection indices (Fig. 5A vs. 5B). This observation is consistent with simulation results and theoretical predictions shown in Fig. 4C, where higher bifurcation probability leads to higher perfection indices.

Next, we asked whether the correlation between bifurcation probabilities and perfection indices also applies to experimental measurements. The measured perfection index for Purkinje cells (0.86, Fig. 2B) is larger than that for class IV neurons (0.70, Fig. 1C). Notably, Purkinje cells have a higher bifurcation probability than class IV cells at lower branch orders, but lower probability at higher orders (Fig. 5C-D). Yet, the perfection indices of Purkinje cells are higher than that of class IV cells. Thus, the bifurcation probability during the early stages of growth decides the perfection index. Moreover, simulations performed with measured bifurcation probabilities vs. branch order (Figs. 5C and 5D) as input were able to reproduce the experimentally observed perfection indices (Fig. S5).

### Quantifying the degree to which dendritic morphologies are stochastic vs. deterministic

The correlation between the bifurcation probability in the Galton-Watson process and the perfection index allows us to relate the stochasticity of a tree to its perfection index. If all the branches with order *k* bifurcate with probability *p*, then the mean number of branches with order *k* + 1 is proportional to *p* and the variance is proportional to *p*(1 – *p*). Because the variance is at a maximum when *p* = 0.5, the percolation threshold, and at a minimum when *p* = 1, we can say that a smaller bifurcation probability and lower perfect index are more stochastic whereas a larger bifurcation probability and higher perfection index are more deterministic. By this measure, the growth rules for mammalian Purkinje cells are less stochastic and more deterministic compared with class IV neurons in insects.

### Tip-support distribution of neuronal branching patterns generated by wiring optimizations also follows a power law

In addition to the Galton-Watson process, we also tested whether other morphogenetic processes give rise to power laws for their tip distribution functions. Cuntz et al. (Cuntz *et al.*, 2008, 2010) proposed a way of constructing branched networks based on an optimal wiring principle using a single parameter, the balancing factor, bf, which weighs the costs of material (total dendrite length) to conduction time (path length to soma). Synthetic trees are grown starting from randomly distributed synaptic target points with *bf* values ranging from a minimum-length tree (*bf* = 0) to a stellate architecture (*bf* = 1). The balancing factor has proved to be an effective parameter to describe a wide variety of neuronal structures from insect dendrites to mammalian neurons (Cuntz *et al.*, 2010; Nanda *et al.*, 2018). A lower balancing factor usually leads to a more branched structure, where the neuron tries to utilize its cytoskeletal resources to the fullest, to fill up a large space. With a higher balancing factor, there are fewer bifurcations as each point of input (for example, synapses or regions of direct sensory reception) demands a relatively direct connection with the soma.

We tested whether trees generated by the balancing-factor process display power laws, we randomly distributed points to mimic synaptic sites and created optimal-wired synthetic trees starting at the center point according to the balancing factor from 0 to 1 in steps of 0.1 (Fig. 6A). We found the tip-support distribution still follows a power law. The perfection index increases monotonically from 0.67 (bf = 0) to 0.81 (bf = 0.9) (Fig. 6B), which falls within the experimentally measured perfection index range. Moreover, it is consistent with the conclusions from previous literature that balancing factors in the range of 0.0 to 0.9 can be used to describe neurons from *Drosophila* sensory neurons to hippocampal granule cells (Cuntz *et al.*, 2010; Nanda *et al.*, 2018).

**Fig. 6.**
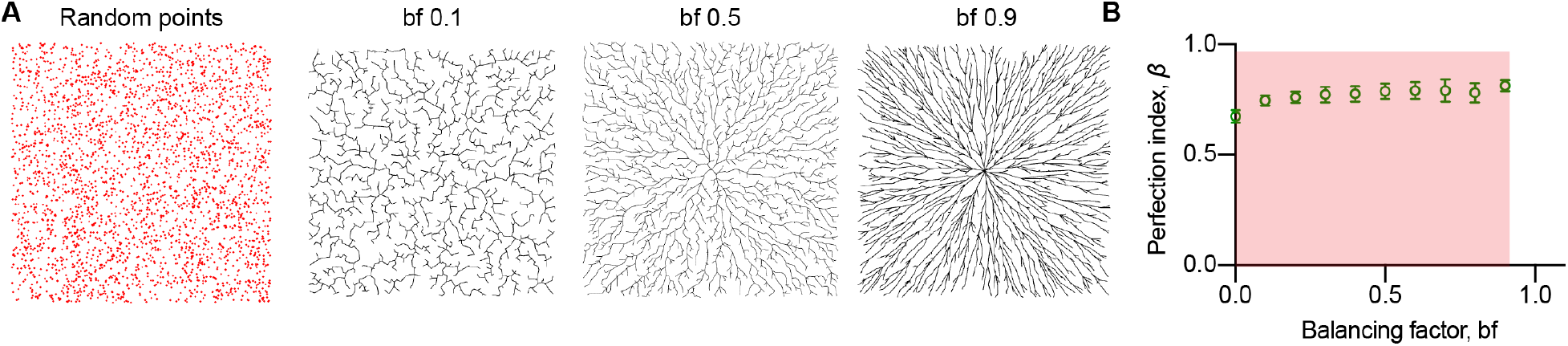
Topological properties of trees generated by different morphogenetic processes. **A.** Example trees grown from randomly distributed points (left panel) from a root at the center. As the balancing factor (bf) increases, the branches become more radial as the trade-off shifts from smaller total branch length to shorter path distances to the center (see text). **B.** Perfection index *β* is plotted as a function of balancing factor. The red shadowed region indicates the region that falls within the experimentally observed range (determined by (mean-2SD, mean+2SD)). Each data point in **B** is averaged from 100 simulations. Error bars are standard deviations.

### Comparison between the perfection index and tree asymmetry

The perfection index is also related to tree asymmetry. A binary partition of *N* terminal branches can be considered as a division of *N* elements (*N* = *N*_1_ + *N*_2_). To indicate the deviation from an equal division, van Pelt et. al. (van Pelt *et al.*, 1992) used a normalized dispersion measure *A*_p_, defined as the division of terminal branches for the partition 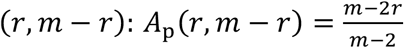 if *m* > 2 with *r* ≤ *m* – *r*. Here *m* – *r* and *r* stand for the number of supported terminal branches for the two bifurcated daughter branches. For a tree of order 1, *A*_p_ (1,1) = 0. The van Pelt tree asymmetry, which is a measure for the asymmetry of the tree, is defined as the weighted mean value of all the *n* – 1 partition asymmetries in the tree with *n* terminal branches. To compare the two measurements, we plotted the tree asymmetry vs. the perfection index for reconstructed neurons from Neuromorpho and Hemibrain datasets (density plot as shown in Fig. 7). Then we generated trees based on Galton-Watson processes and measured the perfection index and van Pelt tree asymmetry simultaneously as shown by the gray dashed curve in Fig. 7. The two quantities are inversely correlated. The perfection index and tree asymmetry estimated from reconstructed neurons tend to cluster towards the gray dashed curve, with average values of 0.8 and 0.5 respectively. Because tree asymmetry has a larger coefficient of variation compared with the perfection index, it might perform better in distinguishing neuronal morphologies of different cell types.

**Fig. 7.**
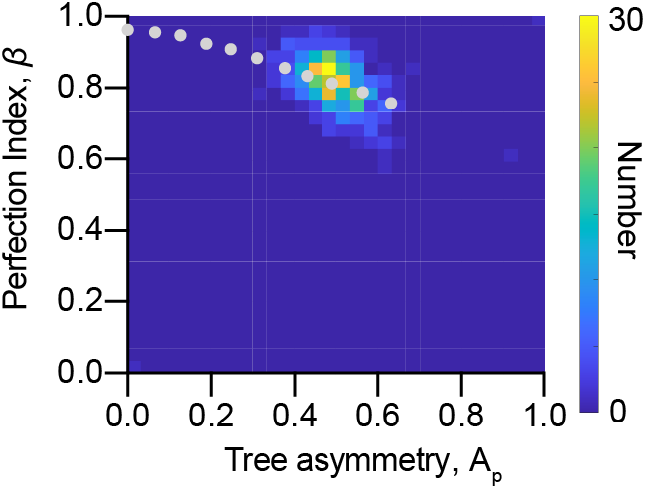
A comparison between the tree asymmetry *A*_p_ and the perfection index *β*. A comparison between the tree asymmetry *A*_p_ and the perfection index *β* measured for 509 cells from NeuroMorpho and hemibrain databases. *A*_p_ =0.50 ± 0.07 and *β* = 0.82 ± 0.07 (mean ± SD, *n* = 509). The dashed curve is obtained from the GW model.

### Randomly adding lateral branches can break the power-law behavior of the tip-support distribution

Next, we asked whether every morphogenetic process gives rise to the power-law behavior of the tip-support distribution. The answer is *no*. In this section, we will elaborate on this observation using both experimental observations and simulations.

There are four classes of sensory neurons that tile the larval body wall (Grueber *et al.*, 2002). The morphology of class III neurons differs from that of class IV neurons, having short branchlets along most of their lengths (Fig. 8A). The tip-support distribution shows two phases (Fig. 8B): a shallower slope for small tip numbers (exponent of −0.84) and a steeper slope for larger tip numbers (exponent of −1.52). When we removed all of the terminal branchlets from class III neuron, thus leaving only what we call the backbone (Fig. S6), we found that the tip-support distribution followed a similar power law (Fig. 8C) as those of class IV neurons with an exponent of −1.5. When we added back branchlets at random locations along the backbone, we recovered the two-phase behavior of the tip-support distribution (Fig. 8B). Thus, class III cells have a backbone with a similar perfection index to class IV cells, with a difference in the tip-support distribution arising from terminal branchlets. This observation suggests that primary branches of class III cells grow by a similar developmental mechanism to class IV cells, i.e., with bifurcation probability above the percolation threshold, followed by the random addition of branchlets, consistent with a recent paper demonstrating that a two-step model is necessary to describe the class III neuron morphology (Stürner *et al.*, 2022).

**Fig. 8.**
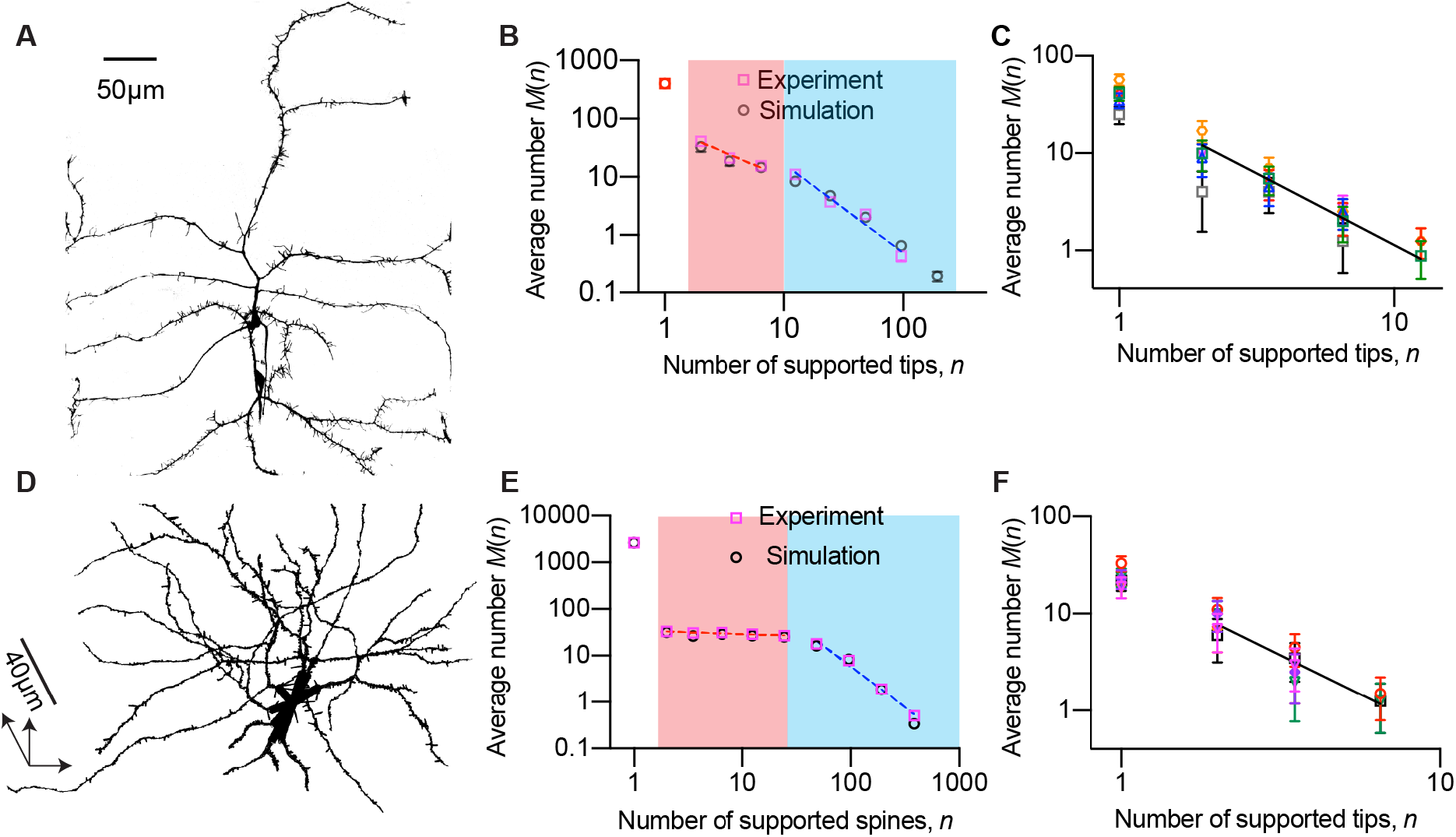
Spines and branchlets lead to deviations from a power law. **A** A GFP-labelled 96hr class III dorsal neuron in the A4 segment imaged by spinning disk confocal microscopy. It has short branchlets along its branches. **B**. The ‘two-phase’ behavior of the tip-support distribution. The red and blue regions have different slopes. **C.** Tip-support distributions for backbones of six different class III neurons (with terminal branches trimmed) in A3-A5 segments from both dorsal and ventral sides fit a power law with slope −1.47. **D** A pyramidal cell from layer 2/3 of mouse visual cortex segmented from the MICrONS electron-microscopy dataset. It has spines along its length. **E** Two-phase behavior of the pyramidal cell with spines is observed. **F.** The tip-support distribution of six trimmed pyramidal cells has a power-law slope −1.61.

Similar to class III neurons, the tip-support distributions of pyramidal cells with spines from in layer 2/3 of the mouse cortex (MICrONs datasets, see Materials and Methods) also deviate from power laws as shown in Fig. 8E. The tip-support distribution shows two phases, a shallow slope with −0.08 and a steeper slope with −1.77. When we removed all spines and measured the tip-support distribution, the power law recovers with an average exponent of 1.6 (Fig. 8F). Note that the pyramidal cells analyzed in Figure 2 lacked spines. Again, adding artificial spines through simulations recovers the experimental observation as shown in Fig. 8E.

## Discussion

In this work, we found that the tip-support distribution, which is a purely topological property of branched networks, follows a power law for most neurons. This distribution is well-characterized by the perfection index, which is half of the absolute value of the power-law exponent and falls within 0.7 to 0.9 with an average of 0.8 over a wide range of neurons. Moreover, this value is invariant to iterative trimming of terminal branches and ablation of internal branches, implying a common branching rule as the neuron grows. We have shown that several morphogenetic processes such as the Galton-Watson process and the optimized wiring procedures produce tip-support distribution that follow power-laws.

To gain a theoretical understanding of the power-law exponent, we mapped the bifurcation process to a percolation transition on the perfect binary tree lattice and demonstrated that this power-law distribution emerges when the bifurcation probability is larger than the percolation threshold, i.e., *p*_c_ = 0.5. When the bifurcation probability is larger than 0.5, one of the daughter branches will probably survive and keep generating offspring over generations. Otherwise, it will become extinct. Thus, the high depth of the branching pattern is the main reason for the emergence of the power-law behavior of the tip-support distribution.

Power-law distributions have been observed for a large range of phenomena in natural, economic, and social systems. Examples range from the number of species in biological taxa (Yule, 1925), the number of cities with a given size (Zipf, 2016), the frequency of earthquakes (Gutenberg and Richter, 1944), the step length in animal search patterns (Viswanathan *et al.*, 1996) and even the spatial distribution of COVID-19 case numbers (Blasius, 2020). We have shown the explanatory power of percolation theory toward the power law that arises from neuronal morphologies, even though they are initially studied in different scientific fields. Such an intriguing connection reemphasizes the nature of the neuron as a physical system. This physics perspective has seen substantial progress in neuroscience. For instance, the idea that the neuronal activity in the cortex might be critical arose from the premise that a critical brain can show the fastest and most flexible adaptation to a rather unpredictable environment (Beggs and Plenz, 2003; Tkačik *et al.*, 2015). We have observed that the power law holds, and the exponent remains roughly invariant under various kinds of operations. This invariance may allow the neuronal morphology to transmit nutrients and information in complex environments, which might themselves have scale-invariant properties such as fractal features. In fact, it has been speculated that scale invariance may confer biological advantages related to the adaptability of response; for example, loss of scale invariance for heartbeat intervals corresponds to a diseased state (Gu *et al.*, 2015).

In addition to rich implications of power-law distributions, we also verified that higher bifurcation probabilities lead to larger power-law slopes and perfection indices from experimental observations, simulations, and theoretical predictions. The stochasticity of binary trees is maximum when the bifurcation probability is 0.5, i.e., at the percolation threshold. On the other hand, the growth rules of binary trees are completely deterministic when the bifurcation probability is 1.0. Thus, the perfection index concept has predictive roles in uncovering the underlying growth mechanisms.

By comparing the perfection index estimated from stochastic bifurcation processes and a wide variety of reconstructed neurons from NeuroMorpho.Org and hemibrain datasets, we found that only simulated trees generated with bifurcation probability near the threshold (between *p*_c_ =0.5 and 0.8) can best describe the observed neuronal branching processes. This observation might imply an optimal choice made by neuronal dendrites, i.e., reducing the amount of cytoplasmic material needed to fill the space by reducing the bifurcation probability while maintaining invariance by growing the dendritic tree after the critical percolation probability. The power-law distribution also holds for synthetic trees generated by a wide range of factors that balance the costs of wiring and path lengths, which has been proven to be effective in describing experimentally reconstructed neuronal structures.

It should be noted that not all neurons have tip-support distributions that satisfy the power-law behavior. Branchlets of class III neurons and spines on pyramidal cells, which can be described by a random lateral branching scheme, can disrupt the power law distribution. The tip-support distribution can therefore be used to distinguish whether more than one branching rule is needed to form neuronal dendrites. It is known that both branchlets of class III neurons and dendritic spines are actin-rich protrusions from neuronal dendrites that dynamically extend and retract (branchlets of class III neurons (Nagel *et al.*, 2012)) or change size and shape (dendritic spines (Hotulainen and Hoogenraad, 2010)) locally. The dendritic shaft or ‘backbone’ of dendrites, on the other hand, is mainly composed of microtubule networks that serve as tracks for long-distance transport (Kapitein and Hoogenraad, 2015). It is possible, therefore, that the different growth mechanisms used to describe protrusions/branchlets (Stürner *et al.*, 2022) and neurites might be attributed to different cytoskeletal components. The characterization scheme that we proposed proved to be an unambiguous criterion for distinguishing the branchlet and spines.

It is worth pointing out that the tip-support distribution should preferentially be used when studying extensive branching patterns, i.e., dendrites with a maximal leaf number larger than 60 since enough datapoints are needed in fitting a power law. Tree asymmetry, on the other hand, can be applied to any tree including ‘smaller trees’. However, different growth schemes, which can be clearly capitulated by the deviation of a power law, cannot be captured by the tree asymmetry concept.

The tip-support distribution we proposed provides a simple and quantitative description of the topology of neuronal morphologies. The power-law slopes of the tip-support distribution have predictive roles in determining neuronal growth mechanisms. This generic property simplifies study of the structure of neurons and thus provides an important additional constraint that must be fulfilled by other methods for generating dendritic trees. It also functions as a method for distinguishing branches with different morphological features.

## Materials and Methods

### *Drosophila melanogaster* strains

*Drosophila melanogaster* larvae were raised on glucose fly food at 25°C. 96hr AEL larvae were used for all analyses. For visualization of dendritic morphologies of class III neurons (ddaF), *nompC-Gal4* (Stock 36361 from Bloomington *Drosophila* Stock Center) was crossed with 10XUAS-mCD8::GFP (32187). The fly strain *ppk-cd4-tdGFP* (a gift from Han Chun (Cornell University)) was used for imaging class IV neurons.

### Spinning Disk Confocal Imaging

Embryos were collected for 2 hrs on apple juice agar plates with a dollop of yeast paste and aged at 25°C in a moist chamber. The plates containing the first batch of embryos were discarded as the dendrite morphology of sensory neurons is less consistent in those animals (Han *et al.*, 2012). Larvae were immobilized individually on agarose pads (thickness 0.3-0.5mm) sandwiched between a slide and a coverslip. The imaging was done using a spinning disk microscope: the Yokogawa CSU-W1 disk (pinhole size 50 μm) built on a fully automated Nikon TI inverted microscope with perfect focus system, an sCMOS camera (Zyla 4.2 plus sCMOS), and running Nikon Elements software. Individual neuron image stacks were acquired with a 60X 1.2 NA water immersion lens with a *z* step size 0.16 μm.

### Data processing method

To reveal the power-law form of the density distribution it is better to plot the density histogram on logarithmic scales. However, the right-hand end of the distribution is noisy because of sampling errors. To deal with it, we vary the width of the bins in the density histogram and normalize the sample counts by the width of the bins they fall in. That is, the number of samples (denoted by 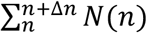) in a bin of width Δ*n* should be divided by Δ*n* to get a count per interval of *n* (i.e., normalized count denoted by *M).* Note that we only consider sample counts larger than 10 to reduce the statistical error. The standard deviation of the normalized count is calculated as: 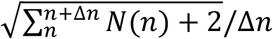. Then the normalized sample count becomes independent of bin width on average and we are free to vary the bin widths as we like. Here we use *logarithmic binning.* We choose a multiplier of 2 and create bins that span the intervals 1.5 to 2.5, 2.5 to 4.5, 4.5 to 8.5 and so forth. The first point N(1), which is just the total number of leaves (or tips), is neglected in the fitting unless the maximal tip number is less than 60 or the total number of normalized sample count is less than 5. The normalized sample counts and the center of the bins are used to plot the results.

### RMA fitting method

Reduced major axis (RMA) regression (Warton *et al.*, 2006; Smith, 2009) is often recommended in allometric scaling analysis for log-transformed data. If it is hard to argue for a cause-effect relationship between X and Y variables, there should be one line describing the pattern of covariation. Neither variable is dependent upon the other, and the biological interpretation of the results should be identical regardless of which variable is on each axis. The RMA slope is the geometrical mean of these two ordinary least square slopes *b_y,x_* and 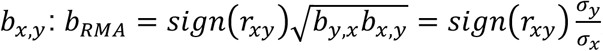, where *r_xy_* stands for the Pearson correlation coefficient between *X* and *Y. σ_y_* and *σ_x_* stand for standard deviations of X and Y variables respectively (detailed description can be found in supplementary information). In our manuscript, the power law exponent *α* is obtained from RMA fitting on log-log transformed data.

### Perfection Index calculation

A function *perfection_tree* is made available as part of the TREES Toolbox (Cuntz *et al.*, 2010) in Matlab.

### Methods for MICrONS dataset analysis

Spine data was recovered from an electron microscopy dataset on layer 2/3 of the mouse visual cortex generated by the MICrONS program. 301 publicly available neuron reconstructions without spines (Turner *et al.*, 2022) were cross-referenced with ~3.2 million automatically identified synapses from the same volume (Dorkenwald *et al.*, 2022). As the synapse dataset is known to contain false-positives, synapses that would imply a spine length of greater than 4 μm were excluded from our analysis.

### Simulating trees with balancing factors

In the functional simulations, the synthetic tree is constrained by the “density profile” of a neuron group and by a balancing factor (bf) (Cuntz *et al.*, 2010) that weighs two demands: the minimization of resources and the minimization of conduction time. Higher bf values correspond to increased importance of conduction time minimization relative to resource minimization and vice versa. The simulation was carried out using the TREES Toolbox package in the MATLAB environment. 3000 random points were generated and synthetic trees starting at the center point according to the balancing factor from 0 to 1 in the step of 0.1 were created for further analyses. For each balancing factor, 100 synthetic trees were created. Note that there exist only two *N(n)* values when balancing factor is set to 1. Thus, the tree perfection index for *bf* = 1 is not calculated in Fig. 6B.

### Data and code availability

All data reported in this paper and all original code are publicly available as the date of publication.

## Appendix

### Derivation of the percolation probability

Given a perfect binary tree with *N*(*N* » 1) branch orders, we associate each edge (branch) with a probability *p*. Each edge in the perfect binary tree will have the probability of either being ‘open’ (with probability *p*) or ‘closed’ (with probability 1 – *p*). An open path in the perfect binary tree is a path where every edge in the path is open. Here we consider the simplest nontrivial case of homogenous edge percolation on the binary tree.

We define the binary survival probability *θ*_p_ as the probability of the simulated tree reaching infinite branch order after branching once. Here, finding *θ*_p_ is equivalent to the probability that a particular vertex *v* is connected to the furthest vertices via paths of open edges. An example of vertices can be found in Fig. A1, where from a root *ψ* a bifurcation results in two vertices *V*_4_ and *V*_2_. It is easy to see that if *p* = 0 then there is no percolation, and that *p = 1* will guarantee percolation.

Let *Z*_2_ be the number of vertices in the binary tree connected to the root and let *Z*_2,*l*_ be the number of vertices connected to the root of branch order *l* from the root. We have 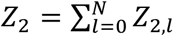. There are 2^*l*^ vertices at branch order *l*. We can label those 2^*l*^ vertices as 1 through 2^*l*^. Then we have 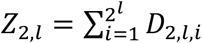, where 1 ≤ *i* ≤ 2^*l*^, *D*_2,*l,i*_ = {*i^th^* vertex at branch order *l* from the root is connected to the root. Thus, *P*(*D*_2,*l,i*_) = *p^l^* and by linearity of expected value we have *E*[*Z*_2,*l*_] = 2^*l*^ *p^l^*. Since *Z*_2,0_ = 1, we obtain:

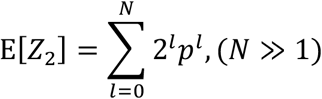

which is finite with 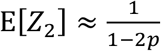 for *p* < 1/2. Hence *θ*_p_ = 0 for *p* < 1/2. For *p* > 1/2, it is also possible to prove that *θ*_p_ is monotonically increasing with *p*.

Note that the probability of the root not percolating is 1-*θ*_p_. If we remove the open edge between root *ψ* and *V*_1_, a new binary tree is formed that is isomorphic to the original binary tree, where vertex *V*_1_ is the new root. Thus, the percolation probability of *V*_1_ on this subtree is equal to *θ*_p_. The probability that the edge between *ψ* and *V*_1_ is open and *V*_1_ percolating on the subtree where *V*_1_ is the root is *pθ*_p_. Therefore, not percolating on the original perfect binary tree with root *ψ*, on the subtree with root *V*_1_ is 1 – *pθ*_p_. Similarly, not percolating on the original perfect tree with root *ψ*, on the subtree with root *V*_2_ is 1 < *pθ*_p_. Since these events are independent of each other, this gives the following equation:

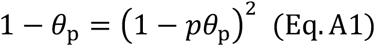

Assume *θ*_p_ ≠ 0. Then we have:

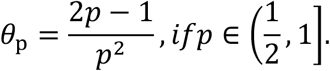

Thus we have:

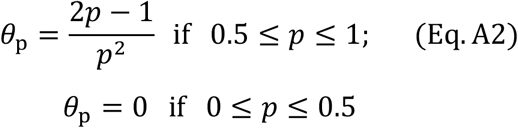

**Fig. A1.**
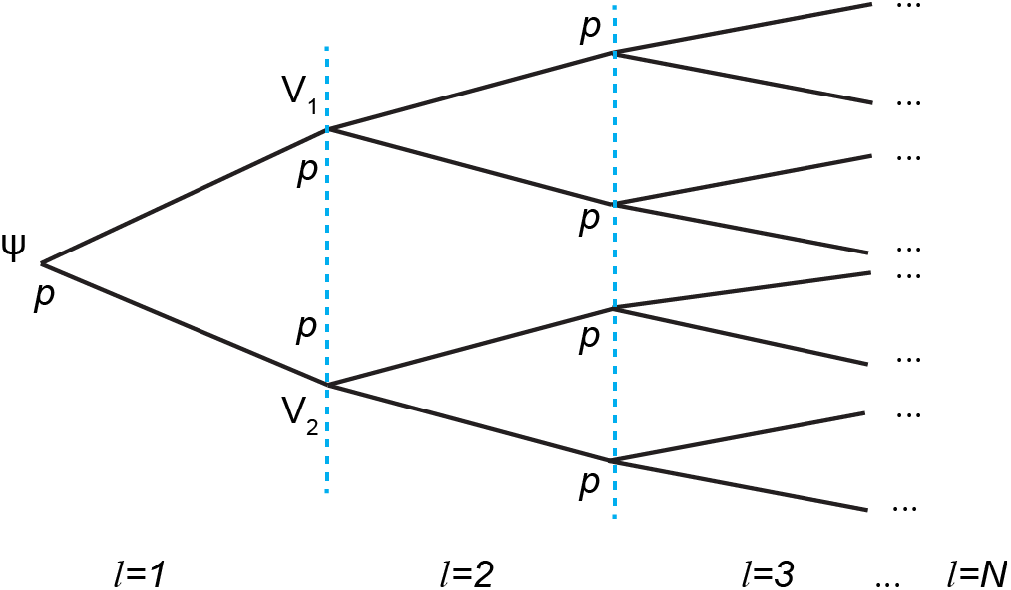
A perfect binary tree with homogeneous edge probabilities. *ψ* stands for root, *V*_1_ and *V*_2_ stand for vertices. *l* stands for branch order. *p* stands for bifurcation probability.

### Power law emerges after the percolation transition

The universal perfection index and power-law distribution observed in various systems motivated us to explore the underlying rules governing the branching in nature. It is generally considered that the appearance of a power law might be an indication that the system operates close to criticality, and it might hint at the presence of a multiplicative stochastic process (Newman, 2005; Sornette, 2006). In this section, we will discuss in detail how the power law originates.

A power-law distribution is also sometimes called a *scale-free distribution* because a power law is the only distribution that is the same on *whatever scale we look at it.* Suppose we have some probability distribution *S*(*x*) for a quantity *x*, and suppose we discover or somehow deduce that it satisfies the property that *S*(*bx*) = *g*(*b*)*S*(*x*) for any *b.* That is, if we increase the scale or units by which we measure *x* by a factor of *b* (coarse grain), the shape of the distribution *S(x)* is unchanged, except for an overall multiplicative constant. Here we will adopt the coarse-grained argument and test whether the observed power law distribution is preserved after one more bifurcation.

Based on our experimental observations, the tip support distribution can be written as *N*(*n*)~*n^-α^*. We can then easily determine how the tip support distribution changes after one more bifurcation on the present binary tree after the percolation transition:

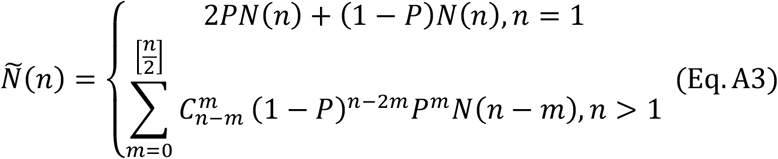

Here [*n*/2] means to take the integer part of *n*/2 and *P* = 2(*p* – *p_c_*), 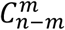 is the binomial coefficient, and 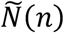 is the updated tip-support distribution after one more bifurcation. If the power law is preserved after one more bifurcation, we should have:

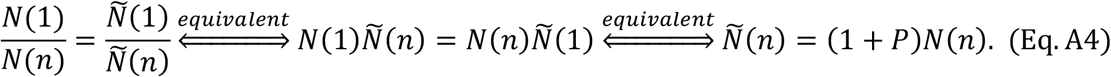

By solving the self-consistent equation Eq. (A4), we find that the power law is preserved after one more bifurcation process as shown in Fig. A2. The exponent *α* gradually increases from 1.7 to 2.0 when the bifurcation probability increases after the percolation point (*p*>0.5, Fig. 4C).

**Fig. A2.**
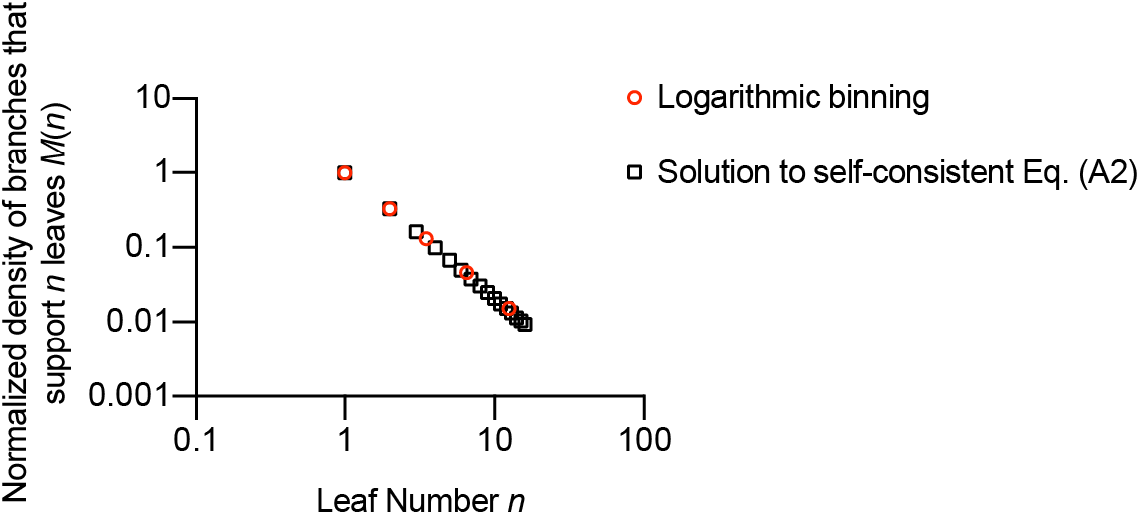
The power law was observed for solved *N*(*n*) of the first 16 leaf numbers after one more bifurcation operation for *p*=0.53 (black square). The data points after logarithmic binning are shown by red circles. The fitted exponent is 1.66.

## Supplementary Information

### Reduced major axis (RMA) linear regression fitting is used based on symmetry consideration

The ordinary least squares (OLS) fitted slope *b* of *Y* on *X* is:

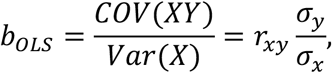

where *COV*(*XY*) stands for the covariance between *X* and *Y. r_xy_* stands for the Pearson correlation coefficient between *X* and *Y.*

A source of concern about OLS is based on the lack of symmetry between the OLS regressions of Y on X and of X on Y. With a single set of data, the biological relationship represented by the equation differs depending upon which variable is assigned to the X axis and which is assigned to Y.

If it is hard to argue for a cause-effect relationship between X and Y variables, there should be one line describing the pattern of covariation. Neither variable is dependent upon the other, and the biological interpretation of the results should be identical regardless of which variable is on each axis.

For a single data set, the difference between the two OLS lines increases as the correlation coefficient decreases. The difference between the two OLS lines is difficult to interpret if they are examined on two separate graphs with reversed axes. However, the X on Y regression can be inverted by fitting Y on X but minimizing the horizontal deviations from the line. The regression of Y on X will be with a slope of: 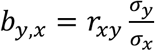. The inverted slope of X on Y will be: 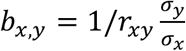. The RMA slope is the Geometrical Mean of these two OLS slopes:

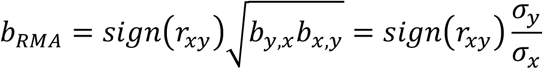

If the axes are inverted for two RMA regressions, the slopes are exact reciprocals of each other, and therefore maintain a single position with respect to the data. Thus, there is only one RMA regression line.

### Definition of perfection index for binary trees in two extreme cases

**Fig. S1.**
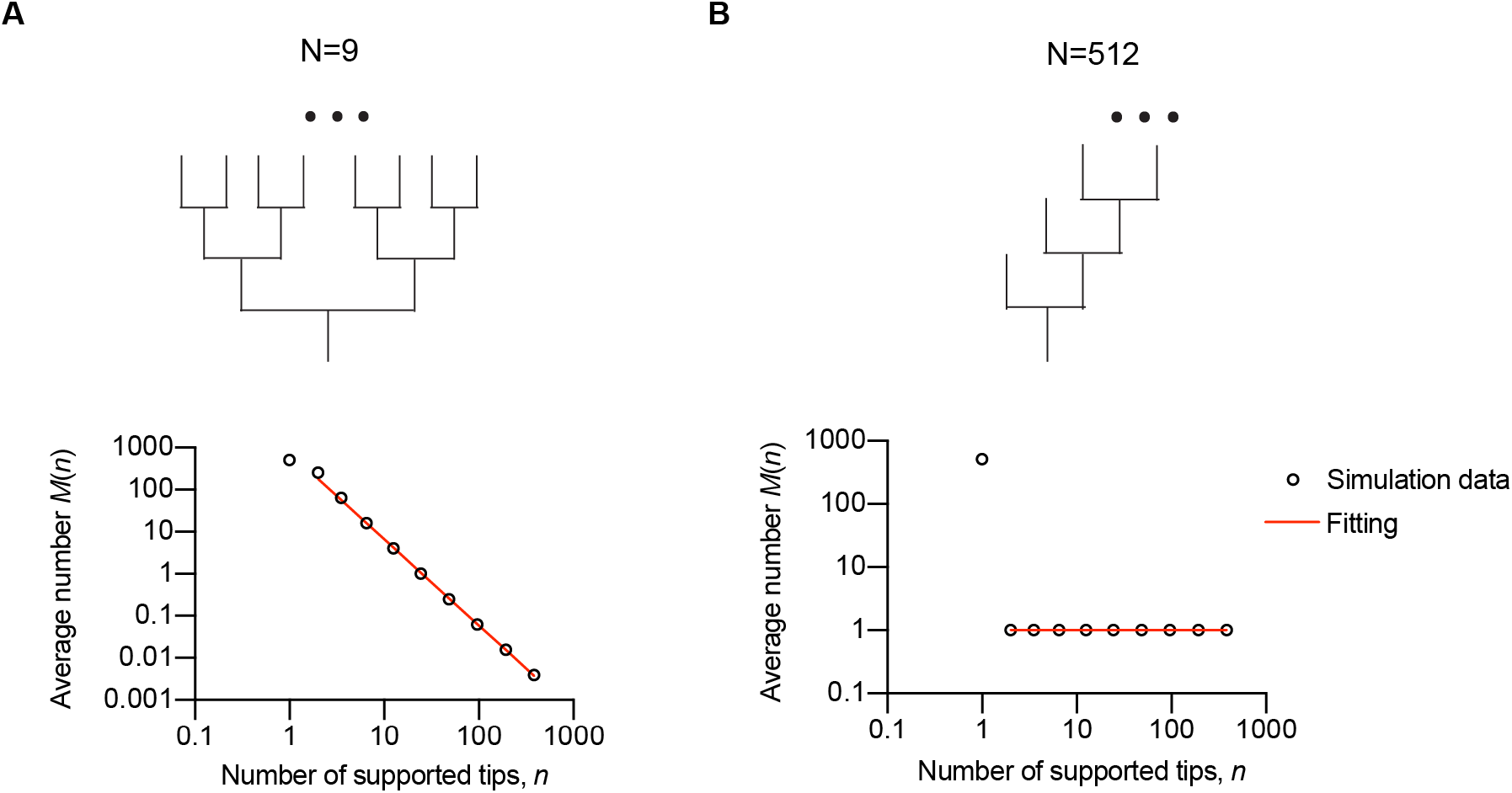
Definition of the tree-perfection index. **A** A perfect tree and **B** A maximally imperfect tree are demonstrated by circular dendrograms. Branches are colored according to the leaf number. In the lower plots, the branch densities are denoted by black circles. The red dashed curves are the RMA regression *y* = *ax^-α^.* Fitted parameters are *α* =2.0 (A) and *α* =0 (B). The first data point is omitted for the reduced major axis (RMA) linear regression fitting. Details of data analysis is described in the Materials and Methods.

### The residual plots for class IV and Purkinje cells confirm the goodness of RMA fitting

**Fig. S2.**
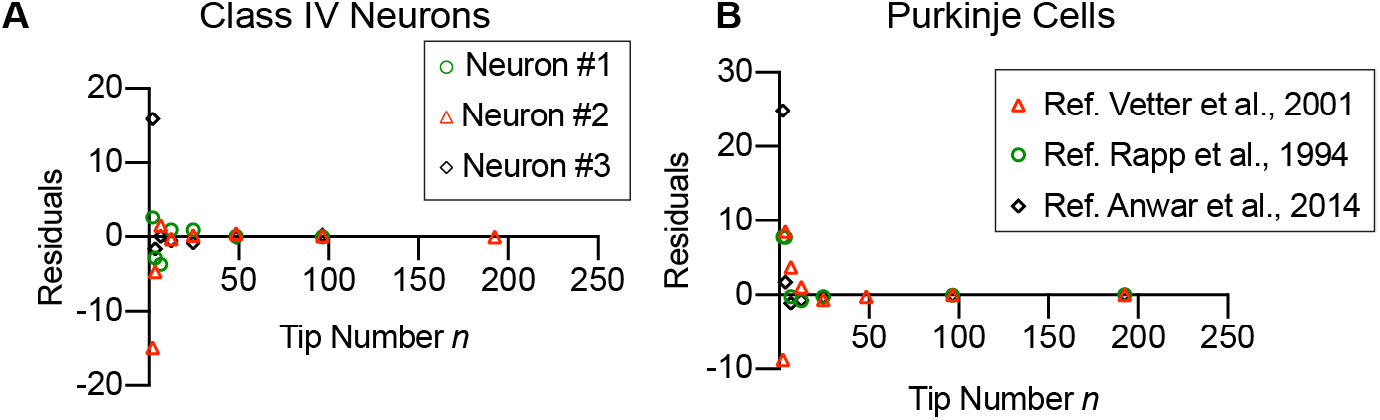
Residual plots of power-law fittings of **A** class IV neurons and **B** Purkinje cells.

### T5 cells in the adult fly central nervous system

**Fig. S3.**
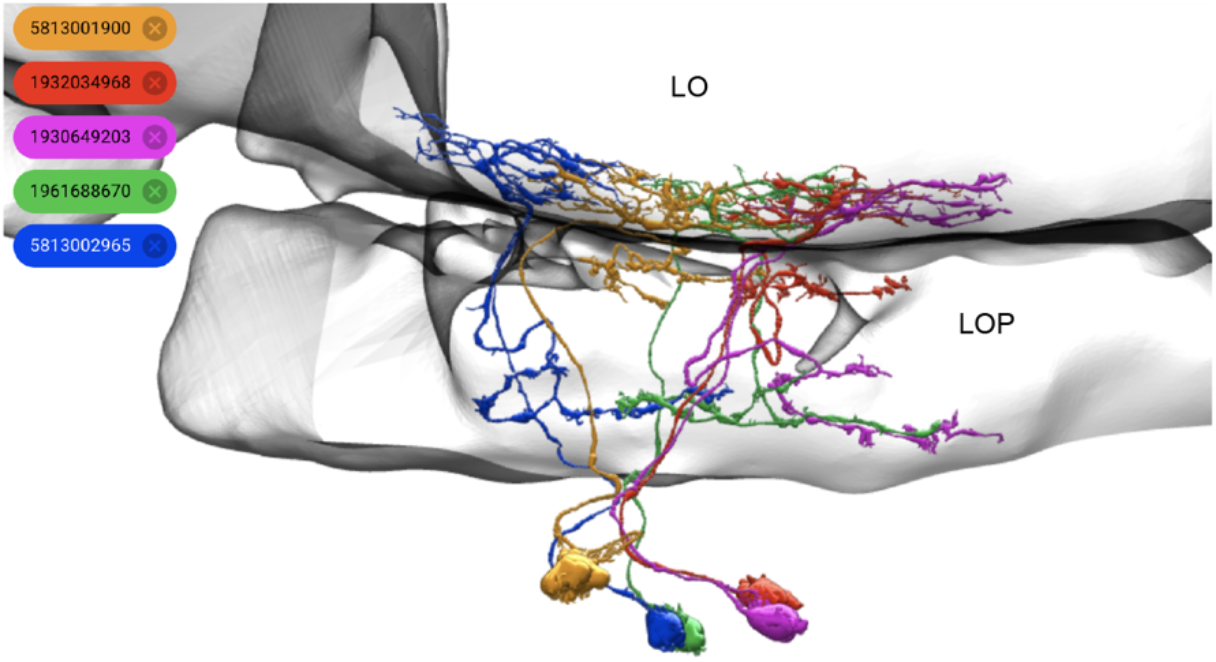
Five examples of T5 cells. Five motion-sensitive T5 cells with dendrites in the lobula (LO) and axons in the lobula plate (LOP) from the Hemibrain dataset. The IDs are indicated at the upper left.

### The power-law distribution of leaf number is observed after the percolation transition for Galton-Watson simulations

**Fig. S4.**
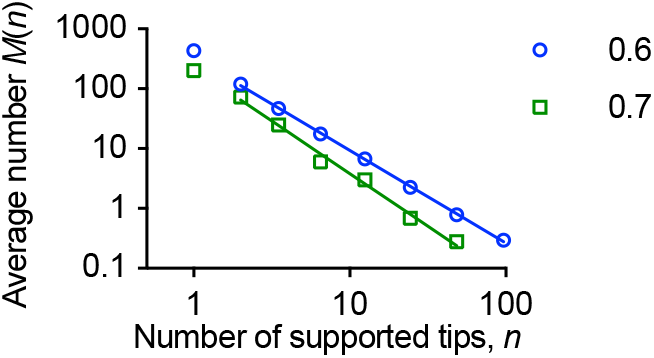
Power-law distributions of simulated trees generated with branching probability 0.6 and 0.7. The perfection indices are 0.78 and 0.88 respectively.

### Galton-Watson simulations based on experimentally measured bifurcation probabilities recapitulate observed perfection indices

In Fig. S5, simulation results using experimentally measured bifurcation probabilities are demonstrated. By using the bifurcation probabilities measured from class IV neurons (Fig. 5C), it is not surprising that the average power-law slope −1.49 from simulations is close to −1.40 which is measured from class IV neurons. Similarly, the average value for Purkinje cells is −1.65, very close to the measured power-law slopes for Purkinje cells, which is −1.72.

**Fig. S5.**
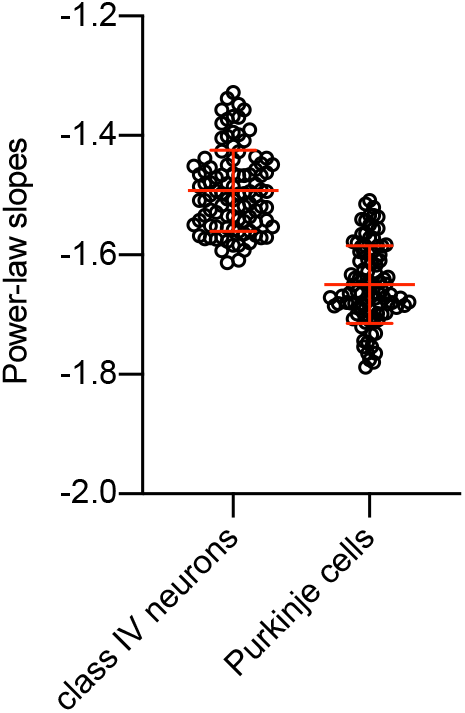
Galton-Watson simulations based on experimentally measured bifurcation probabilities from class IV neurons (Fig. 5C) and Purkinje cells (Fig. 5D). 100 simulations were performed for each case. Powerlaw slopes are −1.49±0.07(*N*=100) for input parameters from class IV neurons; and −1.65±0.06 (mean ± SD, *N*=100) for input parameters from Purkinje cells.

### Simulations show that two-phase behavior can be understood by randomly adding terminal branches to the backbone with a tip-support distribution which follows power law

To further understand the class III neuron morphology, we randomly add the trimmed terminal branches back to the class III neuron backbone as shown in Fig. S6B. The *M*(*n*) vs. *n* relation for the backbone is shown by the green squares in Fig. 8C. The random addition of terminal branches perfectly recapitulates the two-phase behavior measured from experimental data (Fig. 8B).

**Fig. S6.**
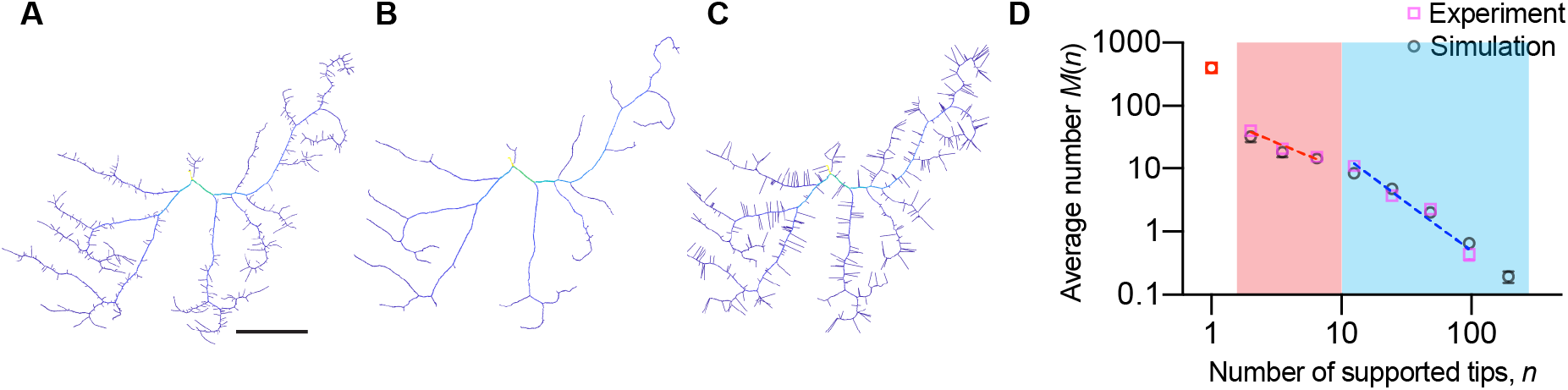
An example of class III neuron. **B** class III neuron with 363 terminal branchlets trimmed. **C** 363 terminal branches with average length 10μm were randomly added back to the backbone as shown in B. Scalebar 50μm. **D** Tip-support distribution for neurons in A and C are plotted together (Fig. 8B).

## ACKNOWLEDGMENTS

The authors thanks Dr. Thomas Torng and Dr. Yin-Wei Kuo for their invaluable feedbacks on this work. This work was supported by NSF GR117147 (J. H.), NIH R01 NS118884 (J. H.), a Burroughs Wellcome Career Award at the Scientific Interface (M.L.), BMBF No. 031L0229 (A. D. B), and Deutsche Forschungsgemeinschaft (DFG, German Research Foundation) No. 467764793, JE 528/10-1 (A. D. B).

## AUTHOR CONTRIBUTIONS

Conceptualization, M.L. and J. H.; Methodology, M.L. and J. H.; Software, A. D. B., M. L. and H. C.; Validation, A. D. B. and H. C.; Formal Analysis, M. L., J. H., A. D. B. and H. C.; Writing, Reviewing and editing, M. L., J. H., A. D. B. and H. C.; Supervision, J. H.

## Reference

Ascoli, GA, and Jeffrey, LK (2000). L-Neuron: a modeling tool for the efficient generation and parsimonious description of dendritic morphology. Neurocomputing 32, 1003–1011.

Anwar, H, Roome, CJ, Nedelescu, H, Chen, W, Kuhn, B, and Schutter, ED (2014). Dendritic diameters affect the spatial variability of intracellular calcium dynamics in computer models. Front Cell Neurosci 8, 168.

Ascoli, GA et al. (2008). Petilla terminology: nomenclature of features of GABAergic interneurons of the cerebral cortex. Nat Rev Neurosci 9, 557–568.

Ascoli, GA, Donohue, DE, and Halavi, M (2007). NeuroMorpho.Org: A central resource for neuronal morphologies. J Neurosci 27, 9247–9251.

Beggs, JM, and Plenz, D (2003). Neuronal avalanches in neocortical circuits. J Neurosci 23, 11167–11177.

Berry, M, and Bradley, PM (1976). The application of network analysis to the study of branching patterns of large dendritic fields. Brain Res 109, 111–132.

Bird, AD, and Cuntz, H (2016). Optimal current transfer in dendrites. Plos Comput Biol 12, e1004897.

Blasius, B (2020). Power-law distribution in the number of confirmed COVID-19 cases. Chaos 30, 093123.

Cajal, S. R. (1995). Histology of the nervous system of man and vertebrates. History of Neuroscience (Oxford Univ Press, New York), 6.

Carlsson, G (2009). Topology and data. B Am Math Soc 46, 255–308.

Caserta, F, Stanley, HE, Eldred, WD, Daccord, G, Hausman, RE, and Nittmann, J (1990). Physical mechanisms underlying neurite outgrowth: A quantitative analysis of neuronal shape. Phys Rev Lett 64, 95–98.

Cuntz, H, Bird, AD, Mittag, M, Beining, M, Schneider, M, Mediavilla, L, Hoffmann, FZ, Deller, T, and Jedlicka, P (2021). A general principle of dendritic constancy: A neuron’s size-and shape-invariant excitability. Neuron 109, 3647-3662.e7.

Cuntz, H, Borst, A, and Segev, I (2007). Optimization principles of dendritic structure. Theor Biol Med Model 4, 21.

Cuntz, H, Forstner, F, Borst, A, and Häusser, M (2010). One rule to grow them all: A general theory of neuronal branching and its practical application. Plos Comput Biol 6, e1000877.

Cuntz, H, Forstner, F, Haag, J, and Borst, A (2008). The Morphological identity of insect dendrites. Plos Comput Biol 4, e1000251.

Cuntz, H, Mathy, A, and Häusser, M (2012). A scaling law derived from optimal dendritic wiring. Proc Natl Acad Sci 109, 11014–11018.

DeFelipe, J et al. (2013). New insights into the classification and nomenclature of cortical GABAergic interneurons. Nat Rev Neurosci 14, 202–216.

Donovan, EJ, Agrawal, A, Liberman, N, Kalai, JI, Chua, NJ, Koslover, EF, and Barnhart, EL (2022). Dendrite architecture determines mitochondrial distribution patterns in vivo. Biorxiv, 2022.07.01.497972.

Dorkenwald, S et al. (2022). Binary and analog variation of synapses between cortical pyramidal neurons. Elife 11, e76120.

Giusti, C, Pastalkova, E, Curto, C, and Itskov, V (2015). Clique topology reveals intrinsic geometric structure in neural correlations. Proc Natl Acad Sci 112, 13455–13460.

Gouwens, NW et al. (2020). Integrated morphoelectric and transcriptomic classification of cortical GABAergic cells. Cell 183, 935–953.e19.

Grueber, WB, Jan, LY, and Jan, YN (2002). Tiling of the Drosophila epidermis by multidendritic sensory neurons. Development 129, 2867–2878.

Grueber, WB, and Sagasti, A (2010). Self-avoidance and tiling: mechanisms of dendrite and axon spacing. Csh Perspect Biol 2, a001750.

Grueber, WB, Ye, B, Yang, C-H, Younger, S, Borden, K, Jan, LY, and Jan, Y-N (2007). Projections of Drosophila multidendritic neurons in the central nervous system: links with peripheral dendrite morphology. Development 134, 55–64.

Gu, C, Coomans, CP, Hu, K, Scheer, FAJL, Stanley, HE, and Meijer, JH (2015). Lack of exercise leads to significant and reversible loss of scale invariance in both aged and young mice. Proc National Acad Sci 112, 2320–2324.

Gutenberg, B, and Richter, CF (1944). Frequency of earthquakes in California*. B Seismol Soc Am 34, 185–188.

Han, C, Wang, D, Soba, P, Zhu, S, Lin, X, Jan, LY, and Jan, Y-N (2012). Integrins regulate repulsion-mediated dendritic patterning of Drosophila sensory neurons by restricting dendrites in a 2D space. Neuron 73, 64–78.

Horak, D, Maletić, S, and Rajković, M (2009). Persistent homology of complex networks. J Statistical Mech Theory Exp 2009, P03034.

Hotulainen, P, and Hoogenraad, CC (2010). Actin in dendritic spines: connecting dynamics to function. J Cell Biol 189, 619–629.

Hwang, RY, Zhong, L, Xu, Y, Johnson, T, Zhang, F, Deisseroth, K, and Tracey, WD (2007). Nociceptive neurons protect Drosophila larvae from parasitoid wasps. Curr Biol 17, 2105–2116.

Jaffe, DB, and Carnevale, NT (1999). Passive normalization of synaptic integration influenced by dendritic architecture. J Neurophysiol 82, 3268–3285.

Jan, Y-N, and Jan, LY (2010). Branching out: mechanisms of dendritic arborization. Nat Rev Neurosci 11, 316–328.

Kanari, L, Dłotko, P, Scolamiero, M, Levi, R, Shillcock, J, Hess, K, and Markram, H (2018). A topological representation of branching neuronal morphologies. Neuroinformatics 16, 3–13.

Kapitein, LC, and Hoogenraad, CC (2015). Building the neuronal microtubule cytoskeleton. Neuron 87, 492–506.

Liao, M, Liang, X, and Howard, J (2021). The narrowing of dendrite branches across nodes follows a well-defined scaling law. Proc Natl Acad Sci 118, e2022395118.

Lin, T-Y, Chen, P-J, Yu, H-H, Hsu, C-P, and Lee, C-H (2021). Extrinsic factors regulating dendritic patterning. Front Cell Neurosci 14, 622808.

Lindenmayer, A (1968). Mathematical models for cellular interactions in development II. Simple and branching filaments with two-sided inputs. J Theor Biol 18, 300–315.

Liu, Z, Wu, M-H, Wang, Q-X, Lin, S-Z, Feng, X-Q, Li, B, and Liang, X (2022). Drosophila mechanical nociceptors preferentially sense localized poking. Elife 11, e76574.

Markram, H et al. (2015). Reconstruction and simulation of neocortical microcircuitry. Cell 163, 456–492.

Markram, H, Toledo-Rodriguez, M, Wang, Y, Gupta, A, Silberberg, G, and Wu, C (2004). Interneurons of the neocortical inhibitory system. Nat Rev Neurosci 5, 793–807.

Marks, WB, and Burke, RE (2007). Simulation of motoneuron morphology in three dimensions. I. Building individual dendritic trees. J Comp Neurol 503, 685–700.

Marshall, W. F. (2020). Scaling of subcellular structures. Annual review of cell and developmental biology, 36, 219–236.

Nagel, J, Delandre, C, Zhang, Y, Förstner, F, Moore, AW, and Tavosanis, G (2012). Fascin controls neuronal class-specific dendrite arbor morphology. Development 139, 2999–3009.

Nanda, S, Das, R, Bhattacharjee, S, Cox, DN, and Ascoli, GA (2018). Morphological determinants of dendritic arborization neurons in Drosophila larva. Brain Struct Funct 223, 1107–1120.

Niven, JE, and Farris, SM (2012). Miniaturization of nervous systems and neurons. Curr Biol 22, R323–R329.

Newman, M (2005). Power laws, Pareto distributions and Zipf’s law. Contemp Phys 46, 323–351.

Pannese, E (2015). Neurocytology, fine structure of neurons, nerve processes, and neuroglial cells. Springer.

Peter, J (1975). Branching processes with biological applications, Wiley.

Rall, W. (1964). Theoretical significance of dendritic trees for neuronal input-output relations. Neural theory and modeling, 73–97.

Rapp, M, Segev, I, and Yarom, Y (1994). Physiology, morphology and detailed passive models of guinea-pig cerebellar Purkinje cells. J Physiology 474, 101–118.

Sartori, F, Hafner, A-S, Karimi, A, Nold, A, Fonkeu, Y, Schuman, EM, and Tchumatchenko, T (2020). Statistical Laws of Protein Motion in Neuronal Dendritic Trees. Cell Reports 33, 108391.

Scheele, CLGJ, Hannezo, E, Muraro, MJ, Zomer, A, Langedijk, NSM, Oudenaarden, A van, Simons, BD, and Rheenen, J van (2017). Identity and dynamics of mammary stem cells during branching morphogenesis. Nature 542, 313–317.

Scheffer, LK et al. (2020). A connectome and analysis of the adult Drosophila central brain. Elife 9, e57443.

Shree, S., Sutradhar, S., Trottier, O., Tu, Y., Liang, X., & Howard, J. (2022). Dynamic instability of dendrite tips generates the highly branched morphologies of sensory neurons. Science Advances, 8(26), eabn0080.

Sizemore, AE, Phillips-Cremins, JE, Ghrist, R, and Bassett, DS (2019). The importance of the whole: Topological data analysis for the network neuroscientist. Netw Neurosci 3, 656–673.

Smith, RJ (2009). Use and misuse of the reduced major axis for line-fitting. Am J Phys Anthropol 140, 476–486.

Snider, J, Pillai, A, and Stevens, CF (2010). A universal property of axonal and dendritic arbors. Neuron 66, 45–56.

Song, W, Onishi, M, Jan, LY, and Jan, YN (2007). Peripheral multidendritic sensory neurons are necessary for rhythmic locomotion behavior in Drosophila larvae. Proc Natl Acad Sci 104, 5199–5204.

Sornette, D (2006). Critical phenomena in natural sciences, chaos, fractals, self-organization and disorder: concepts and tools. Springer Series Syne.

Sterling, P, and Laughlin, S (2015). Principles of Neural Design, MIT Press.

Strahler, A. N. (1952). Hypsometric (area-altitude) analysis of erosional topography. Gsa Bulletin 63, 1117–1142.

Stürner, T, Castro, AF, Philipps, M, Cuntz, H, and Tavosanis, G (2022). The branching code: A model of actin-driven dendrite arborization. Cell Reports 39, 110746.

Tkačik, G, Mora, T, Marre, O, Amodei, D, Palmer, SE, Berry, MJ, and Bialek, W (2015). Thermodynamics and signatures of criticality in a network of neurons. Proc Natl Acad Sci 112, 11508–11513.

Turner, NL et al. (2022). Reconstruction of neocortex: Organelles, compartments, cells, circuits, and activity. Cell 185, 1082–1100.e24.

Van Pelt, J., Uylings, H. B., Verwer, R. W., Pentney, R. J., & Woldenberg, M. J. (1992). Tree asymmetry—a sensitive and practical measure for binary topological trees. B Math Biol 54, 759–784.

Van Pelt, J., van Ooyen, A., & Uylings, H. B. (2000). Modeling dendritic geometry and the development of nerve connections. In Computational Neuroscience. CRC Press. (200–229)

Vetter, P, Roth, A, and Häusser, M (2001). Propagation of action potentials in dendrites depends on dendritic morphology. J Neurophysiol 85, 926–937.

Viswanathan, GM, Afanasyev, V, Buldyrev, SV, Murphy, EJ, Prince, PA, and Stanley, HE (1996). Lévy flight search patterns of wandering albatrosses. Nature 381, 413–415.

Vormberg, A, Effenberger, F, Muellerleile, J, and Cuntz, H (2017). Universal features of dendrites through centripetal branch ordering. Plos Comput Biol 13, e1005615.

Warton, DI, Wright, IJ, Falster, DS, and Westoby, M (2006). Bivariate line-fitting methods for allometry. Biol Rev 81, 259–291.

Wen, Q, and Chklovskii, DB (2008). A cost–benefit analysis of neuronal morphology. J Neurophysiol 99, 2320–2328.

Wen, Q, Stepanyants, A, Elston, GN, Grosberg, AY, and Chklovskii, DB (2009). Maximization of the connectivity repertoire as a statistical principle governing the shapes of dendritic arbors. Proc Natl Acad Sci 106, 12536–12541.

Werginz, P, Raghuram, V, and Fried, SI (2020). The relationship between morphological properties and thresholds to extracellular electric stimulation in RGCs. J Neural Eng 17, 045015.

Williams, AH, O’Donnell, C, Sejnowski, TJ, and O’Leary, T (2016). Dendritic trafficking faces physiologically critical speed-precision tradeoffs. Elife 5, e20556.

Yule, GU (1925). II.—A mathematical theory of evolution, based on the conclusions of Dr. J. C. Willis, F. R. S. Philosophical Transactions Royal Soc Lond Ser B Contain Pap Biological Character 213, 21–87.

Zipf, GK (2016). Human behavior and the principle of least effort: An introduction to human ecology, Ravenio Books.

